# Development of a sandwich ELISA for the detection of bovine A1 beta-casein

**DOI:** 10.1101/2025.10.17.682492

**Authors:** Ayumi Watanabe, Tomoe Kobayashi, Anna Okamoto, Daiki Oka, Tomohiro Noguchi, Ren Ozawa, Koumei Shirasuna, Makoto Matsuyama, Takashi Kuramoto

**Author notes:** Correspondence to Takashi Kuramoto, Ph.D., Laboratory of Animal Nutrition, Department of Animal Science, Faculty of Agriculture, Tokyo, University of Agriculture, 1737 Funako, Atsugi, Kanagawa 243-0034, Japan; Makoto Matsuyama, Ph.D., Division of Molecular Genetics, Shigei Medical Research Institute, 2117-18 Yamada, Minami-ku, Okayama-shi, Okayama, 701-0202, Japan.

## Abstract

The genetic variant A2 beta-casein is associated with fewer digestive and absorption issues compared to A1 beta-casein, leading to increased global demand for A2 milk. However, contamination with the A1 variant during collection, transportation, or sterilization of A2 milk poses a risk, necessitating a verification test to ensure A2 milk does not contain A1 beta-casein. We developed an A1-specific monoclonal antibody (mAb) and a general mAb that reacts with both A1 and A2 variants using the iliac lymph node method. A sandwich ELISA was created using the general mAb as the capture antibody and the A1-specific mAb as the detection antibody to identify A1 beta-casein in milk. This ELISA successfully detected A1 beta-casein in raw and pasteurized A2 milk, including ultra-high temperature treated milk. The test identified A1 beta-casein when the A1 spike in A2 milk exceeded 1% in volume, indicating its capability to detect contamination from one A1A1 cow in a herd of one hundred A2A2 cows. The developed A1 beta-casein ELISA is suitable for high-throughput analysis and can be valuable for monitoring A1 beta-casein contamination in commercially produced A2 milk.

## Introduction

The proteins in milk consist of 80% casein and 20% whey proteins. Beta-casein accounts for approximately 30–45% of the casein [1,2] and is classified into A1 and A2; A1 has the histidine residue at the 67th amino acid position, whereas A2 has the proline residue at that position [3]. The amino acid substitution is derived from a single nucleotide substitution from A to C at the codon (CAT to CCT) in exon 7 of the bovine beta-casein 2 gene.

Beta-casomorphins are opioid-like peptides that are released during enzymatic digestion of beta-casein in the intestine [4]. Owing to the amino acid substitution, beta-casomorphin-7 (BCM7) is released from A1 but not from A2 beta-casein [5,6]. Studies suggest that bovine BCM7 is a risk factor for cardiovascular disease, type 1 diabetes, sudden infant death syndrome, and neurological disorders such as autism and schizophrenia [5,7,8]. BCM7 is also involved in intestinal physiology, such as mucin release at the intestinal lumen [9], activation of immune-inflammatory cells [10], and IgA secretion [11]. These findings suggest that A1 beta-casein can have adverse effects on human health, and efforts have thus been made to produce A1-free A2 milk.

Most of the scientific work related to A2 milk has focused on milk intolerance and gastrointestinal (GI) symptoms. Studies on the effects of milk have examined serum biomarkers, stool frequency and consistency, GI transit time, constipation, stomach and bowel inflammation, and fecal conditions in healthy people as well as in patients with lactose intolerance. Although a systematic review concluded that there is insufficient human-based evidence from clinical trials to prove the adverse digestive health effects of A1 compared with A2 milk [12], more recent studies found that A2 milk decreases GI symptoms such as gastrointestinal inflammation [13], rapid gastric emptying [14], and prolonged digestive discomfort in lactose intolerant people [14,15]. Even in healthy toddlers, A2 milk improved overall digestive comfort and GI-related symptoms [16]. These findings suggest that A2 milk can improve the symptoms similar to lactose intolerance.

The potential positive impact of A2 milk on human health has promoted the production of A2 milk by farmers around the world. The production of herds consisting of only A2A2 cows has been the aim of farmers in various countries such as New Zealand [17], Brazil [18], Mexico [19] Turkey [20], Italy [3], and Poland [21]. Because of the increase in the global demand for A2 milk, parts of Japanese dairy farmers are focusing on producing A2 milk, and they expect that A2 milk and its products (e.g., yogurt and infant formula) will promote the sustainable development of the dairy industry by providing the high-prized dairy products to the Japanese market.

One challenge in producing and distributing A2 milk is to prove that A2 milk is free from A1 beta-casein. Although A2 milk does not contain A1 beta-casein, there is a potential risk of contamination during the collection, transportation, and sterilization of A2 milk. Because such contamination may lead to unintentional falsification, a verification system for A2 milk is required. Indeed, a falsification was found in Austria, in which four of the five milk samples analyzed contained A1 beta-casein [22].

To prevent falsification, several methods to detect A1 beta-casein in milk have been developed using isoelectric focusing [22], high-performance liquid chromatography, and mass spectrometry [23]. Although each of these methods has specific advantages, they are all time-consuming and labor intensive. For routine application, a solid-phase enzyme-linked immunosorbent assay (ELISA) system is superior because it saves time and labor and offers high-throughput analysis.

ELISA can detect and quantify target proteins in a sample solution, but requires an antibody that is specific to the target molecule. For A1 beta-casein, a few IgY antibodies are available, among which one is used in a commercially available ELISA kit (Biosensis, Pty Ltd., Thebarton, Australia) and another one was developed in-house [24]. Both are polyclonal antibodies (pAbs). Meanwhile, monoclonal antibody (mAb) has advantages over pAb. The hybridoma allows for the continuous production of mAb without the need for repeated immunization [25]. Thus, we tried to produce a new mAb specific to A1 beta-casein and used it to develop an ELISA system.

In the present study, we used the iliac lymph node method [26] to produce beta-casein specific and A1-specific mAbs. We developed a sandwich ELISA system using these antibodies and showed that it detects an A1 spike in A2 milk at a 1% spike level in both raw and processed milk, providing a method to test the purity of A2 milk.

## Materials & Methods

### Preparation of control samples

A1A1 and A2A2 milk samples for screening hybridoma were provided by a dairy farm in Fujinomiya, Shizuoka, Japan. These milk samples were collected from cows with known A1A1 (n=1) and A2A2 (n=1) beta-casein genotypes. The milk samples were kept on ice until they arrived at the laboratory from the dairy farm. Fifty milliliters of fresh milk was stirred while being kept on ice. Milk fat aggregated was filtered through cotton wool. We then centrifuged the milk samples at 7200×g at 4°C for 15 minutes [27] and filtered the supernatant through 5A filter paper (Toyo Roshi Kaisha, Ltd., Tokyo, Japan). The defatted milk samples were diluted 1:100 with phosphate buffered saline (PBS), and stored at −80°C.

### Milk samples for evaluation of ELISA

Individual cow milk samples were provided by a dairy farm in Furano, Hokkaido, Japan. Raw milk was collected from healthy cows with known beta-casein genotypes (n = 67) and stored at 4□°C until arrival at the laboratory. Upon arrival, the samples were stored at −80□°C until further experiments. To evaluate the specificity of the antibody against A1 beta-casein, defatted milk samples from 22 cows (A1A1: n = 11; A2A2: n = 11) were used. To assess the performance of the sandwich ELISA using raw milk samples, 35 samples were tested (A1A1: n = 4; A1A2: n = 13; A2A2: n = 18). To determine the detection limit of the A1 beta-casein ELISA, 10 samples (A1A1: n = 5; A2A2: n = 5) were subjected to three different processing conditions: raw, ultra-high temperature (UHT) treatment, and pasteurization.

### Ultra-high temperature processing and pasteurization

Pasteurization was performed by heating at 63°C for 30 min. Ultra-high temperature (UHT) processing of the raw milk was performed by heating at 121°C for 2 s using a plate heat exchanger (HAS-CH-360, IWAI KIKAI KOGYO CO., LTD. Tokyo, Japan) The sterilized samples were diluted 1:100 and stored at −80°C.

### Production of monoclonal antibodies

Mouse and rat monoclonal antibodies (mAbs) were produced using the iliac lymph node method [26]. Briefly, the A1-specific peptide (67His) with the sequence 67His: NH_2_-CFPGPIH^67^NSLP-COOH (Eurofins Genomics, Tokyo, Japan) was conjugated to keyhole limpet hemocyanin (KLH), and an antigen emulsion was injected into BALB/cJ mice (The Jackson Laboratory Japan, Yokohama, Japan) or WKY/NCrl rats (CLEA Japan, Tokyo, Japan). The treated mice and rats were euthanized on day 21 after the injection, and lymphocytes were fused with SP2/0-Ag14 myeloma cells. After the cell fusion, culture supernatants were screened to select positive clones, which reacted with the 67His but not with the 67Pro (NH_2_-CFPGPIP^67^NSLP-COOH), by solid-phase ELISA. Among positive clones, we isolated a clone that reacted to A1 milk but not A2 milk.

The mAb reacting to both A1 and A2 was generated in a similar manner using a KLH-conjugated synthetic peptide with the sequence NH_2_-CESLSSSEESITRINKKIEK-COOH as the antigen. This peptide corresponds to the residues from 29 to 47 of the bovine beta-casein [28]

### Western blotting

Milk proteins were purified from A1A1 and A2A2 milk samples. To remove fat, raw milk samples were stirred on ice and centrifuged at 7200 × g for 15□min at 4□°C, following the method described by Kato et al. [27]. The resulting supernatant was filtered through 5A filter paper (Toyo Roshi Kaisha, Ltd., Tokyo, Japan). Defatted milk was then freeze-dried and stored at −20□°C until use.

To denature the milk proteins, 0.2□g of freeze-dried milk powder was dissolved in 20□mL of PBS (pH 7.4) containing 0.5% SDS and 2% 2-mercaptoethanol. The mixture was thoroughly stirred overnight using a rotary shaker. The extract was then centrifuged at 10,000 × g for 30□min at room temperature, and the supernatant was filtered through a 0.8□μm membrane filter (AGC TECHNO GLASS Co., Ltd., Shizuoka, Japan). Bovine beta-casein was purchased from Sigma-Aldrich (C6905, St. Louis, MO, USA) and dissolved in 0.1□M Tris-HCl buffer (pH 8.0). Purified milk proteins (4.2□μg) from A1A1 and A2A2 samples, along with bovine beta-casein, were subjected to electrophoresis on 15% SDS-PAGE. The gel was stained with Coomassie Brilliant Blue (CBB Stain One, Nacalai Tesque, Inc., Kyoto, Japan). Following electrophoresis, proteins were transferred onto polyvinylidene fluoride membranes. To block nonspecific binding, membranes were incubated with 5% BSA in blocking buffer for 1□h at room temperature. The membranes were then incubated at 4□°C for 24□h with either a mouse mAb raised against an A1 beta-casein-specific peptide, or a rat mAb raised against a peptide common to both A1 and A2 beta-casein, each diluted 1:3000 in blocking buffer. After washing, membranes were incubated for 1□h at room temperature with an HRP-conjugated goat anti-mouse IgG secondary antibody (Fortis Life Sciences, Boston, MA, USA) diluted 1:3000, or a goat anti-rat IgG secondary antibody (Abcam, Cambridge, UK) diluted 1:10,000, both in blocking buffer. Signal development was performed using Clarify Western ECL substrate (Bio-Rad Laboratories, Hercules, CA, USA), and signals were detected with an ImageQuant LAS 4000 image analyzer (GE Healthcare Japan Corporation, Tokyo, Japan).

### Sandwich ELISA

The mAb that reacted with both A1 and A2 was used as the capture antibody, and the A1-specific mAb was used as the detection antibody in the sandwich ELISA. Briefly, 1 mg of the capture antibody in 100 μL of PBS was added to each well of the Nunc-Immuno Plate (Thermo Fisher Scientific, Tokyo, Japan) and incubated overnight at 37°C. Then, 200 μL of blocking buffer (1% BSA/PBS) was added and incubated at 37°C for 1 h. Next, 100 μL of milk sample diluted to 1:10000 with PBS was added and incubated at 37°C for 2 h. The detection antibody (0.1 μg / 100 μL in 1% BSA/PBS) was added and incubated at 37°C for 2 h. Goat anti–mouse IgG2a-HRP enzyme–labeled secondary antibody (Bethyl Laboratories, Montgomery, TX, USA) was diluted to 1:20000 with 1% BSA/PBS, and 100 μL of the diluted secondary antibody was added to the sample and incubated at room temperature for 1 h. Finally, 150 μL of developing solution consisting of 0.04% *o*-phenylenediamine (FUJIFILM Wako Pure Chemical Corporation, Osaka, Japan) and 0.04% H_2_O_2_ were added and incubated at room temperature for 30–60 min. The reaction was stopped by addition of 50 µL of 3M H_2_SO_4_. Optical density (OD) was measured using a plate reader (Tecan Japan, Kawasaki, Japan) at a wavelength of 492 nm. Before adding each reagent except for H_2_SO_4_, the solution in wells was discarded and the wells were washed 2–3 times with PBS.

### Evaluation of bovine A1 beta-casein ELISA

The detection limit for A1 beta-casein was determined using a discriminative point approach [29]. We prepared 5 sets of A1A1-A2A2 test mixtures which contained 1%, 2%, 3%, 10%, 25%, and 50% A1A1 milk spikes, respectively. Briefly, we collected raw milk samples from five A1A1 cows and five A2A2 cows, respectively. We combined the individual A2A2 milk equally to prepare the A2A2 milk mixture. We then added individual A1A1 milk with the A2A2 mixture to make five sets of A1A1-A2A2 test mixtures. These mixtures consisted of six different ratios, in which A1A1 milk was contained at 1%, 2%, 3%, 10%, 25%, and 50%, respectively. Along with 100% A1A1 and 100% A2A2 milks, these mixed samples were subjected to the ELISA test in triplicate. Test sample mixtures of pasteurized and UHT milk were prepared in the same manner.

Cut-off value is calculated using the following equation:

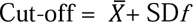

where *X̄* is the mean of OD values of negative controls, SD is the standard deviation, and *f* is the multiplier which is derived from the critical values for a one-tailed *t*-distribution and obtained from the number of negative controls and the confidence level [29]. In the present study, we obtained the *f* value at 95% confidence level.

When the OD value was lower than the cut-off, the mixture was considered negative for A1 beta-casein. When the OD value was equal to or higher than the cut-off, the mixture was considered positive for A1 beta-casein.

## Results

### The monoclonal antibody detected A1 beta-casein but not A2 beta-casein

We obtained a monoclonal antibody that reacted with the 67His (A1) peptide, but not with the 67Pro (A2) peptide. To examine the specificity of the antibody for A1 beta-casein, we performed sandwich ELISA using defatted A1A1 (n=11) and A2A2 (n=11) milk. The OD from PBS (n = 3) was 0.17 ± 0.0005. As the multiplier factor (*f*) was 3.37 at 95% of confidence level, the cut-off value was calculated as 0.172. The mean OD of A1A1 milk was 1.56 ± 0.29 and the OD values of the A1A1 milk samples were all higher than the cut-off value. Consistently, the mean OD of A2A2 milk was 0.13 ± 0.04 and the OD values of the A2A2 milk samples were all lower than the cut-off value (**Fig. 1**). These results indicated that the monoclonal antibody generated detected A1 beta-casein but not A2 beta-casein in the defatted milk.

**Figure 1.**
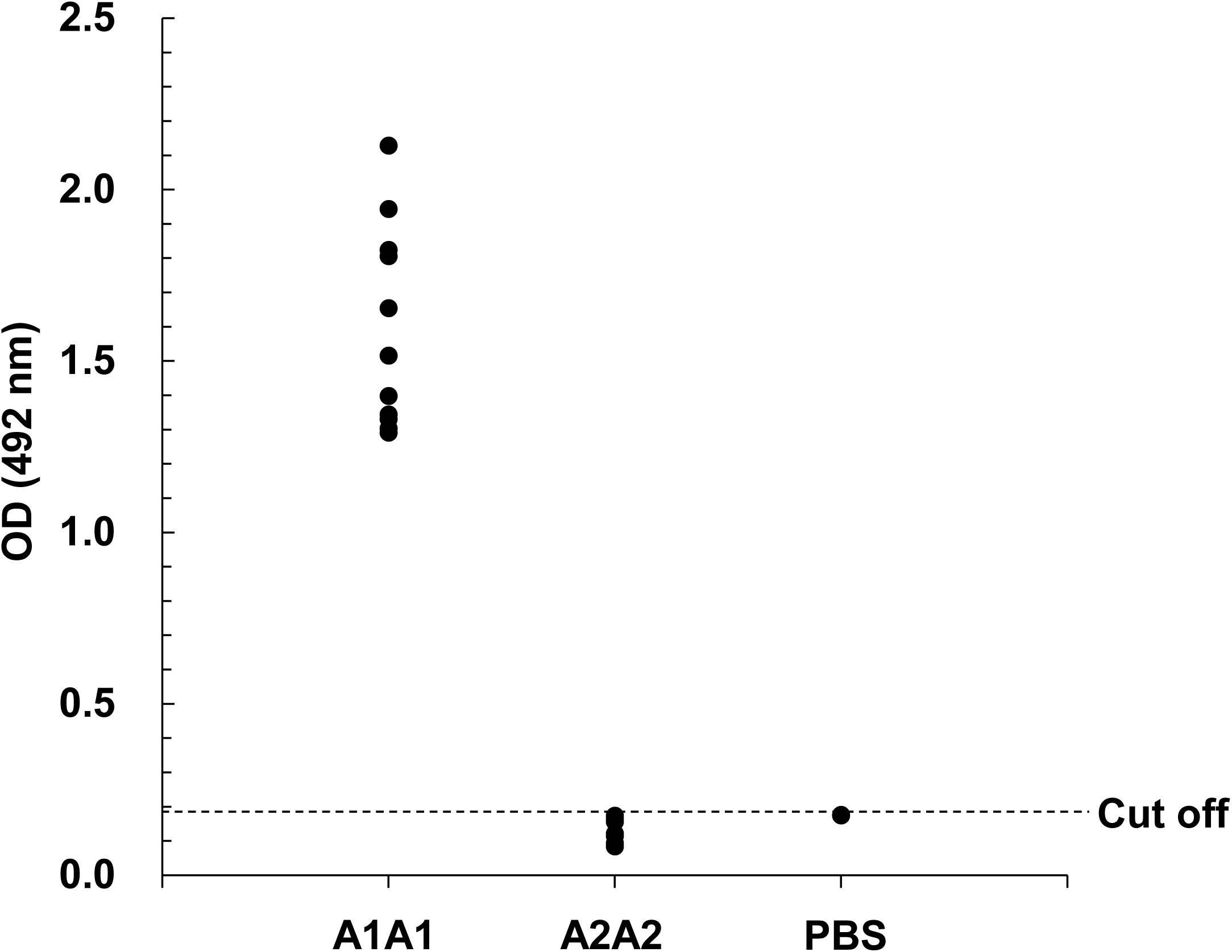
Distribution of OD values obtained from ELISA for defatted milk samples. OD values were obtained from A1A1 (n = 11) and A2A2 (n = 11) samples. The horizontal dashed line represents the cut-off value (0.17) calculated from the OD of PBS.

To further confirm the specificity of the mAbs, we performed Western blot analysis using milk proteins purified from A1A1 and A2A2 milk. Purified bovine beta-casein (C6905, Sigma-Aldrich) was included as a positive control. The mAb raised against a peptide common to both A1 and A2 beta-casein detected protein bands in the 21–31□kDa range from both A1A1 and A2A2 milk proteins (**Fig. 2B**). In contrast, the mAb raised against an A1-specific peptide detected a 21–31□kDa band only in the A1A1 milk protein and not in the A2A2 sample (**Fig. 2C**). These results further support that the mAb specific to A1 beta-casein selectively detects A1 but not A2 beta-casein.

**Figure 2.**
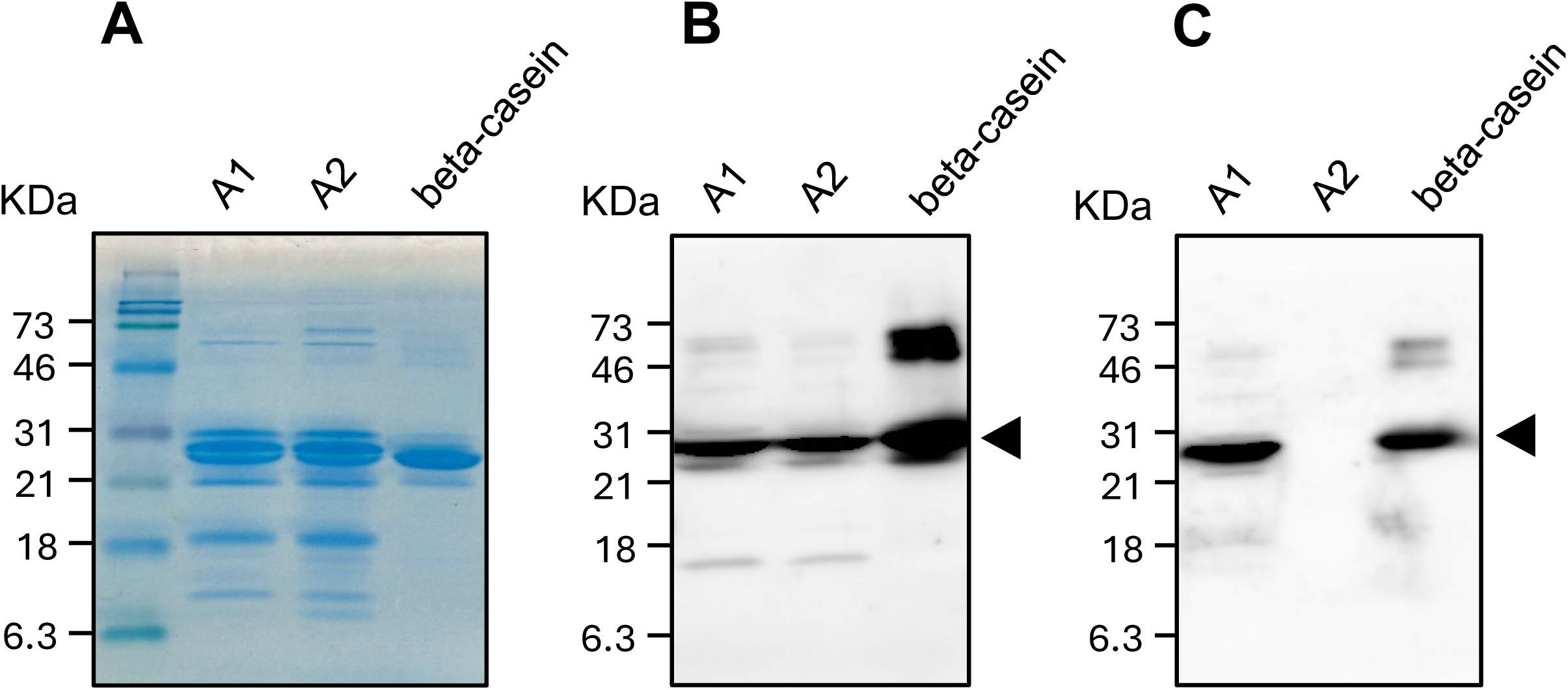
Western blot analysis of milk protein purified from A1A1 and A2A2 milk. A, Proteins separated on SDS-PAGE gel were stained with Coomassie Brilliant Blue. B, Western blotting analysis with rat mAb raised from the peptide shared with A1 and A2 beta-casein. This mAb detected 21–31 kDa bands from both proteins obtained from A1A1 and A2A2 milk as well as from the purified bovine beta-casein (arrowhead). Bands found in the purified beta-casein (46–73 kDa) might be derived from self-association of the beta-casein[36]. C, Western blotting analysis with mouse mAb raised from the peptide specific to A1 beta-casein. The mAb detected 21–31 kDa bands from milk protein obtained from A1A1 and the purified beta-casein (arrowhead) but not from A2A2 milk. A1, denatured whole milk protein from A1A1 milk. A2, denatured whole milk protein from A2A2 milk. beta-casein, purified bovine beta-casein purchased from Sigma Aldrich (C6905).

### Evaluation of the sandwich ELISA using raw milk samples

We found that the sandwich ELISA was applicable to defatted milk. Given that the ELISA was designed to test bulk milk, a mixed raw milk that was collected from different cows and stored in a bulk tank of dairy farm, we first tried to detect A1 beta-casein in raw milk samples. We performed the sandwich ELISA using raw A1A1 (n = 4), A1A2 (n = 13), and A2A2 (n = 18) milk samples. The OD from PBS (n = 3) was 0.19 ± 0.003 and the cut-off value was 0.200.

The OD values of the A1A1 and A1A2 raw milk samples were all higher than the cut-off value, and the mean OD values of A1A1 and A1A2 milk were 1.19 ± 0.04 and 1.04 ± 0.11, respectively. Consistently, the OD values of A2A2 raw milk were all lower than the cut-off value, and the mean OD of A2A2 milk was 0.17 ± 0.03 (**Fig. 3**).

**Figure 3.**
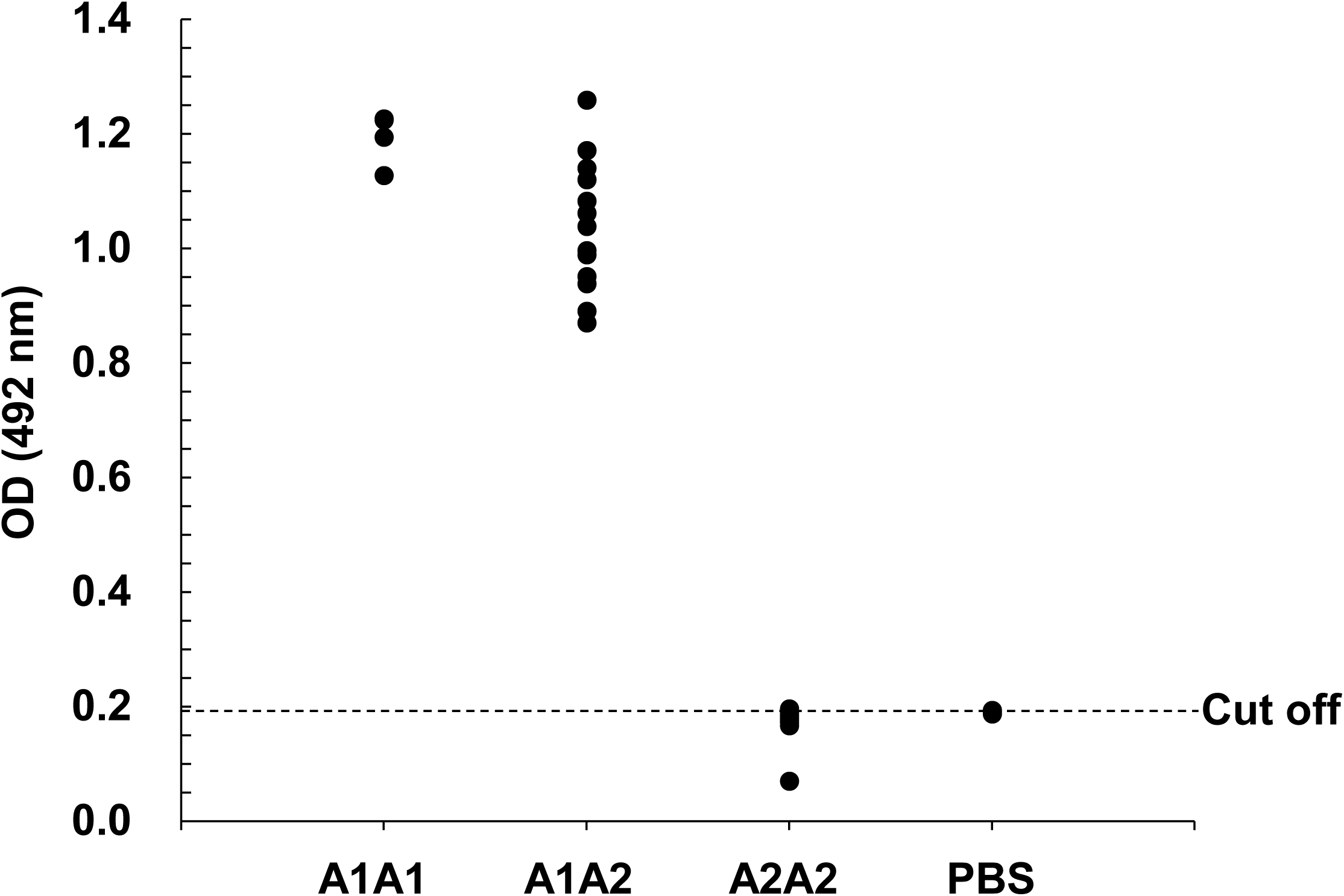
Distribution of OD values obtained from ELISA for raw milk samples. OD values were obtained from A1A1 (n = 4), A1A2 (n = 13), and A2A2 (n = 18) samples. The horizontal dashed line represents the cut-off value (0.20) calculated from the OD of PBS.

Next, we performed the ELISA using samples containing a mixture of A1A1 and A2A2 milk to determine whether an A1A1 milk spike could be detected in A2A2 milk. The OD from negative control (pure A2 milk) was 0.15 ± 0.01 for raw milk. As the factor (f) was 1.82 at a 95% confidence level, the cut-off value was calculated as 0.16. As shown in **Table 1**, the OD values obtained from test mixtures with 50%, 25%, 10%, 3%, 2%, and 1% spikes were higher than the cut-off value, which was calculated to be 0.16, indicating that these samples were positive for A1 beta-casein. These findings suggest that the sandwich ELISA was able to detect A1 beta-casein in raw milk with an A1A1 milk spike of 1% or more in A2A2 milk.

**Table 1.** Mean OD values obtained from raw, pasteurized, and UHT milk containing different spike levels of A1.

| % A1<br>in A2 | Raw |  | Pasteurized |  | UHT |  |
| --- | --- | --- | --- | --- | --- | --- |
|  | OD <sup>1</sup> | CI <sup>2</sup> | OD | CI | OD | CI |
| 100 | 1.10±0.06 | 1.07–1.13 | 1.25±0.13 | 1.19–1.32 | 0.84±0.10 | 0.80–0.89 |
| 50 | 0.81±0.08 | 0.77–0.85 | 0.93±0.09 | 0.88–0.97 | 0.62±0.04 | 0.60–0.64 |
| 25 | 0.69±0.07 | 0.65–0.73 | 0.72±0.05 | 0.70–0.75 | 0.49±0.03 | 0.47–0.50 |
| 10 | 0.44±0.05 | 0.42–0.47 | 0.46±0.04 | 0.44–0.48 | 0.33±0.03 | 0.31–0.35 |
| 3 | 0.29±0.04 | 0.27–0.32 | 0.32±0.08 | 0.28–0.37 | 0.26±0.05 | 0.23–0.28 |
| 2 | 0.23±0.03 | 0.21–0.24 | 0.23±0.03 | 0.21–0.24 | 0.19±0.03 | 0.17–0.20 |
| 1 | 0.18±0.01 | 0.18–0.19 | 0.19±0.02 | 0.18–0.20 | 0.16±0.01 | 0.15–0.16 |
| 0 | 0.15±0.01 | 0.14–0.15 | 0.15±0.01 | 0.14–0.15 | 0.12±0.004 | 0.11–0.12 |

1, Mean ± SD of OD values from 15 measurements.
2, Confidence interval with 95% confidence level

### Evaluation of the sandwich ELISA using sterilized milk samples

We found that the sandwich ELISA could detect A1 beta-casein in raw milk. Because the ELISA is also expected to be used for sterilized milk, we attempted to detect A1 beta-casein in sterilized milk samples. The OD from negative control (pure A2 milk) was 0.15 ± 0.01 for pasteurized milk, and 0.12±0.004 for UHT milk. As the factor (f) was 1.82 at a 95% confidence level, the cut-off value was calculated as 0.16 for pasteurized milk and 0.12 for UHT milk. As shown in **Table 1**, the 50%, 25%, 10%, 3%, 2%, and 1% pasteurized mixtures showed larger OD values than the cut-off value, which was calculated to be 0.16, indicating that these pasteurized samples were positive for A1 beta-casein. In the case of UHT milk, the cut-off value was 0.12, and the 50%, 25%, 10%, 3%, 2%, and 1% UHT mixtures were also positive for A1 beta-casein. These findings suggest that the sandwich ELISA was able to detect A1 beta-casein in sterilized milk with a 1% spike of A1A1 milk in A2A2 milk.

## Discussion

In the present study, we developed a mAb specific to the bovine A1 beta-casein. We also developed a mAb that could react with both A1 and A2 beta-casein. The mAb against the beta-casein was used as the capture antibody and the A1-specific antibody was used as the detection antibody to develop a sandwich ELISA for detecting A1 beta-casein in milk.

To develop mAbs, we initially screened hybridomas using the peptides employed for immunization. ELISA using defatted milk samples demonstrated that the mAb raised against the peptide shared by both A1 and A2 beta-casein reacted with both A1 and A2 milk, whereas the mAb raised against the A1-specific peptide reacted with A1 milk but not with A2 milk.

To verify that these mAbs were targeting beta-casein itself, we performed Western blot analysis. The mAb raised against the shared peptide detected a 21–31□kDa band in both A1 and A2 milk proteins, which corresponded to the band observed for purified beta-casein (Fig. 2B). In contrast, the mAb raised against the A1-specific peptide detected the band in A1 milk protein, but not in A2 milk protein (Fig.2C). These results indicate that both mAbs recognize beta-casein, and that the mAb raised against the A1-specific peptide selectively detects A1 beta-casein.

The sandwich ELISA detected A1 casein when A1A1 milk was present at a 1% spike level in A2A2 milk, suggesting that the detection level of the ELISA was 1% in volume. We also showed that the sandwich ELISA was applicable to sterilized milk, indicating that the ELISA can be used to monitor A1 beta-casein contamination in A2 milk before and after milk processing.

We evaluated the detection level of the ELISA by determining the lowest percentage of A1 milk in A2 milk at which OD values were significantly larger than those of the negative control (pure A2 milk). To identify significant differences, we used a threshold value, or cut-off. The cut-off value was calculated from the OD values of the negative controls, and if the OD values of the samples were larger than the cut-off, the samples were considered to contain A1. In other words, we considered that the ELISA could detect A1 at the given concentration in the sample.

There are several methods for setting the cut-off value, such as defining it as two to three times the mean OD of the background measurement or the negative control. However, these approaches are arbitrary and lack statistical justification. Therefore, we determined the cut-off based on the method described by Frey et al. [29]. This method aims to establish an appropriate detection limit by using Student’s *t*-distribution and thereby calculates a statistically justified cut-off. This approach has been applied to set a threshold for determining positive and negative results in ELISA-based detection of *E*. *histolytica* infection [30]. Similarly, de Jesus et al. applied this method to establish a statistically significant threshold in ELISA-based detection of A1 beta-casein [24].

Here, we used pure A2 milk as the negative control and set the cut-off with a 95% confidence level using the formula described by Frey et al. [29]. We found that the OD values within the 95% confidence interval of the 1% spike of A1 raw milk, as well as sterilized milk, did not overlap with the cut-off values. Thus, we considered the detection limit of the A1 beta-casein ELISA to be 1%, suggesting that the ELISA is capable of detecting contamination from one A1A1 cow in a herd of one hundred A2A2 cows.

There are several anti-beta-casein antibodies produced. A murine mAb 1C3 was raised by using purified beta-casein as an antigen. Radioimmunoassay showed that the 1C3 bound the peptide from 193 to 202 of beta-casein [31,32]. Oudshoorn et al. produced three murine mAbs using purified beta-casein as an antigen and identified their epitope at residues 105–193 [33]. Senocq et al. raised 21 murine mAbs against purified beta-casein as an antigen and characterized epitopes. The mAbs recognized 14 distinct epitopes clustered in six discrete determinants (4–8, 14–24, 33–48, 49–91, 178–183, 184–209) [34]. Our mAbs reacting both A1 and A2 and the A1 solely were produced using the synthetized peptides with residues 29–47 and 61–71, respectively. Thus, the epitopes of the mAbs were very likely to differ from the published mAbs reacting the bovine beta-casein.

There are two anti-A1 antibodies available; one is a commercially available polyclonal anti-A1-IgY (Biosensis Pty Ltd, Australia), and the other one is a polyclonal anti–A1-IgY that was generated in-house by de Jesus et al.[24]. Biosensis developed a sandwich ELISA kit that includes the IgY as the detection antibody. The kit can detect spikes of bovine A1A1 milk in goat and camel milk at 50%, 25%, and 5% spike level. Although the kit can detect A1 milk in A2 milk at a 5% spike level, data on the detection of lower spike levels are not available. Moreover, the Biosensis kit requires the dilution of milk samples with NaOH, which is a harmful reagent for humans.

The bovine A1 beta-casein ELISA system developed in this study can detect A1 casein at a 1% spike level in raw as well as sterilized milk and does not require NaOH treatment, and the results indicate that it is more reliable and safer than the conventional ELISA systems.

The North American A2 milk market is expected to increase exponentially by 2029 because of consumers’ preference for milk with superior properties compared with regular milk. Similarly, the European market is anticipated to demonstrate robust growth driven by research and development activities in the dairy sector [35]. In Japan, some dairy farmers have established herds of A2A2 cows and are providing A2 milk to the market. A verification system to demonstrate the absence of contamination of A1 beta-casein in A2 milk is thus desirable. This verification consists of genotyping tests for cows and protein tests for the milk product. The A1 beta-casein ELISA developed in this study is expected to contribute to the verification of A2 milk. Given the growing interest in A2 milk worldwide, the ELISA would be used globally and contribute to the development of A2 dairy products for commercial purposes, as well being useful for quality inspection of A2 dairy products by the government.

In summary, we developed a new A1 beta-casein ELISA method capable of detecting an A1 spike in A2 milk at a 1% spike level. The results of the analysis suggest that the detection level of the ELISA is sufficient for use in the routine monitoring of A2 milk.

## Acknowledgements

The authors are grateful to the dairy farms that provided milk samples. This study was supported in part by the Japan A2 Milk Association. Watanabe A. is a recipient of Support for Pioneering Research initiated by the Next Generation program of the Japan Science and Technology Agency.

## Note

The monoclonal antibodies against bovine A1 beta-casein are protected by a patent (No. PCT/JP2023/034219).

